# Mathematical models of tumor volume dynamics in response to radiotherapy

**DOI:** 10.1101/2022.04.07.487525

**Authors:** Nuverah Mohsin, Heiko Enderling, Renee Brady-Nicholls, Mohammad U. Zahid

## Abstract

From the beginning of the usage of radiotherapy (RT) for cancer treatment, mathematical modeling has been integral to understanding radiobiology and for designing treatment approaches and schedules. There has been extensive modeling of response to RT with the inclusion of various degrees of biological complexity. Here we focus on models of tumor volume dynamics. There has been much discussion on the implications of different models of tumor growth, and it is just important to consider the implications of selecting different models for response to RT. In this study, we compare three models of tumor volume dynamics: (1) exponential growth with RT directly reducing tumor volume, (2) logistic growth with direct tumor volume reduction, and (3) logistic growth with RT reducing the tumor carrying capacity. For all three models, we: performed parameter sensitivity and identifiability analyses; investigated the impact of the parameter sensitivity on the tumor volume trajectories; and examined the rates of change in tumor volume (ΔV/Δt) during and RT treatment course. The parameter identifiability and sensitivity analyses revealed the interdependence of the different model parameters and may inform parameter calibration in any further usage of these models. In examining the ΔV/Δt trends, we coined a new metric – the point of maximum reduction of tumor volume (MRV) – to quantify the magnitude and timing of the expected largest impact of RT during a treatment course. Ultimately, the results of these analyses help us to better understand the implications of model selection while simultaneously generating many hypotheses about the underlying radiobiology that need to be tested on time-resolved measurements of tumor volume from appropriate pre-clinical or clinical data. The answers to these questions and more detailed study of these and similar models of tumor volume dynamics may enable more appropriate model selection on a disease-site or patient-by-patient basis.

## 1. Introduction

Radiotherapy (RT) is the most common oncologic agent with more than 50% of all cancer patients receiving radiation as part of their cancer treatment (Delaney et al. 2005). From the beginning of the usage of RT, mathematical modeling has been integral to designing treatment approaches and schedules. Nearly all of radiobiology and RT treatment schemes are built on fitting clonogenic survival assay curves to the linear-quadratic (LQ) model, which assumes that there is a linear and a quadratic component involved in cell death by RT (Fowler 1984, 1989, 1990; Brenner 2008). There has been extensive mathematical modeling work done in the field of RT for a range of purposes: understanding radiobiology (Barendsen 1982; Brenner et al. 1998; Alfonso and Berk 2019); predicting response to particular radiation doses/schedules (Enderling et al. 2010; Prokopiou et al. 2015; Jeong et al. 2017; Sunassee et al. 2019; Hormuth et al. 2021; Liu et al. 2021; Zahid et al. 2021a, b); personalizing dose, dose distributions and fractionation schedules (Leder et al. 2014; López Alfonso et al. 2014; Alfonso et al. 2018; Zahid et al. 2021b, c). These modeling efforts have varied in the degree of biological complexity included, from bottom-up models of DNA damage & repair (Sachs et al. 1992), vascular damage (Powathil et al. 2013; Rodríguez-Barbeito et al. 2019), oxygenation (Jeong et al. 2013; Scott et al. 2016; Lewin et al. 2018, 2020; Grimes 2020), immune interactions (Poleszczuk et al. 2016; Serre et al. 2016; Poleszczuk and Enderling 2018), sub-populations of competing tumor cells(Paczkowski et al. 2021), etc. to more coarse-grained models of gross tumor volume (Rockne et al. 2010; Prokopiou et al. 2015; Watanabe et al. 2016; Cho et al. 2020; Zahid et al. 2021a, b). We are particularly focused on top-down models of tumor volume dynamics that may be more tractably calibrated to tumor volumes abstracted from regularly acquired clinical images, such as CT or MRI scans.

There has been much discussion on the implications of different models of tumor growth (Gerlee 2013). Similarly, it is important to consider the implications of selecting different models for response to RT. Commonly, RT mechanism of action is assumed to be direct tumor cell kill, as governed by the LQ model. Recently, we have introduced the concept of modeling the effect of RT as a reduction of carrying capacity in a logistic growth model of tumor growth (Zahid et al. 2021b). This is premised on the fact that RT treatment fields encompass the entire tumor and do not selectively kill tumor cells – rather they also effect the tumor microenvironment, which includes microvasculature, immune cells, and other stromal cells. The assumption is that each RT fraction reduces the capacity of the local tissue to support a tumor. This leads to either reduced tumor growth, if the carrying capacity remains higher than the tumor volume, or tumor volume reduction, otherwise. Here, we compare three models of tumor volume dynamics and compare the implications of the model assumptions in terms of the effects on rates of change in tumor volume, parameter sensitivity and parameter identifiability.

## 2. Methods and Models

### 2.1 Models of Tumor Growth and Response to RT

We studied three models of tumor volume dynamics that consist of all the combinations of 2 models of tumor growth (exponential and logistic) and 2 modes of response to RT (direct tumor volume reduction and carrying capacity reduction).

#### 2.1.1 Exponential Growth with Direct Tumor Volume Reduction (EXP+DVR) Model

Exponential tumor volme growth is modeled as:

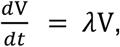

where V(t) is the tumor volume [cc] at time t and *λ* is the intrinsic volumetric tumor growth rate [day^-1^]. In this model, tumor response to each radiation fraction is modeled with an exponential instantaneous reduction in tumor volume:

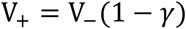

where V_+_ [cc] is the tumor volume post radiation fraction, *V*_−_ [cc] is the tumor volume prior to radiation fraction, and *γ* is the tumor volume reduction parameter. *γ* is defined by the linear-quadratic (LQ) dose response model: 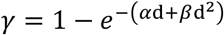, where *α* [Gy^-1^] and *β* [Gy^-2^] describe the cell’s radiosensitivity, commonly described by the fraction 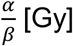 (McMahon 2018). Dose [Gy], d, is tumor radiation exposure for each fraction.

#### 2.1.2 Logistic Growth with Direct Tumor Volume Reduction (LOG+DVR) Model

Logistic tumor volume growth is modeled as:

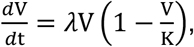

where V(t) is the tumor volume [cc] at time t, *λ* is the intrinsic tumor growth rate [day^-1^], and K [cc] is the tumor carrying capacity, which is calculated as follows:

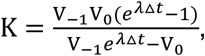

V_−1_ and V_0_ are two temporally spaced pretreatment measurements of tumor volume at the time of radiotherapy planning and prior to the first radiation fraction, respectively; and *Δt* is the time difference between the V_−1_ and V_0_ measurements. We have previously introduced the proliferation saturation index, 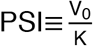, to describe the proliferative capacity of a tumor (Prokopiou et al. 2015). PSI is defined between 0 and 1, where a tumor with PSI = 0 exhibits pure exponential growth and a tumor with PSI = 1 does not grow. In alignment with the Norton-Simon hypothesis that the rate of tumor response to therapy is proportional to the tumor growth rate (Norton and Simon 1977; Norton et al. 1977; Simon and Norton 2006), we modeled tumor response to each radiation fraction with an instantaneous tumor volume reduction modulated by the instantaneous tumor volume to carrying capacity ratio:

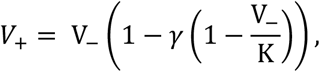

where V_+_ [cc] is the tumor volume post radiation fraction and *V*_−_ [cc] is the tumor volume prior to radiation fraction. Similar to the EXP+DVR model, *γ* is the cell death parameter defined by the LQ dose response model, with parameters of d [Gy] and 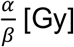.

#### 2.1.3 Logistic Growth with Carrying Capacity Reduction (LOG+CCR) Model

The LOG+CCR model is the same as the LOG+DVR model, except that the tumor carrying capacity is not a constant because the impact of radiotherapy is modeled as an instantaneous carrying capacity reduction:

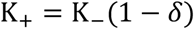

where K_+_ is the tumor carrying capacity post radiation fraction, K_-_ is the tumor carrying capacity prior to radiation fraction, and *δ* is the carrying capacity reduction fraction defined between 0 and 1. When *δ* = 0, the carrying capacity remains unchanged and when *δ* = 1, the carrying capacity is reduced to 0. In this model, a radiation fraction dose only results in tumor volume reduction if *K*_+_ < V_−_. This allows for tumor growth early in treatment when *K*_$_ > *V*_−_. These three models are summarized and compared in Table 1.

**Table 1.** Description of radiotherapy mathematical models

|  | Exponential Growth with Direct Tumor Volume Reduction (EXP+DVR) | Logistic Growth with Direct Tumor Volume Reduction (LOG+DVR) | Logistic Growth with Carrying Capacity Reduction (LOG+CCR) |
| --- | --- | --- | --- |
| Volume Dynamics | $\frac{dV}{dt} = \lambda V$ | $\frac{dV}{dt} = \lambda V \left(1 - \frac{V}{K}\right)$ | |
| Effect of Radiation | $V_+ = V_-(1-\gamma)$ | $V_+ = V_- \left(1 - \gamma \left(1 - \frac{V_-}{K}\right)\right)$ | $K_+ = K_-(1-\delta)$ |
| Varied Parameters | $\lambda, \alpha$ | $\lambda, \alpha, \text{PSI}^{**}$ | $\lambda, \delta, \text{PSI}^{**}$ |
| Fixed Parameters | $d^*, \frac{\alpha^*}{\beta}$ | $d^*, \frac{\alpha^*}{\beta}$ | |
| Initial conditions | $V_0$ | $V_0$ | $V_0, K_0$ |

\* $\gamma = 1 - e^{-(\alpha d + \beta d^2)}$ , where dose, $d$ , and $\frac{\alpha}{\beta}$ are the biological parameters in the EXP+DVR and LOG+DVR models.
\*\* Proliferation saturation index, $\text{PSI} = \frac{V_0}{K_0}$ , where $V_0$ and $K_0$ are the tumor volume and carrying capacity prior to initiation of radiotherapy, respectively. While $K = K_0$ in the LOG+DVR model, $K$ varies with each radiation fraction in the LOG+CCR model.

### 2.2 Model Simulations

All simulations were performed using custom MATLAB scripts. Tumor volume changes were simulated with a time resolution of 1 hour for a total time of 8 weeks. All tumor volume model simulations begin with two weeks of pretreatment tumor growth followed by a six-week RT treatment schedule. Pre-treatment tumor growth occurs between the time points *t*_−1_ and *t*_0_, the time at radiation planning and prior to the first fraction of radiotherapy, respectively. The treatment schedule consists of radiation fractions delivered once daily Monday through Friday for six weeks to mimic standard fractionated RT as delivered in many clinical settings.

Example simulations for all three models, with parameters selected to fit tumor volumes of a patient with head and neck cancer treated at Moffitt Cancer Center with 2 Gy standard fractions, can be seen in Figure 1. The plots illustrate the capacity of all three models to describe observed volume dynamics under a real treatment regimen. Notably, 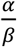 is set to 10 Gy whenever the LQ model is utilized.

**Figure 1.**
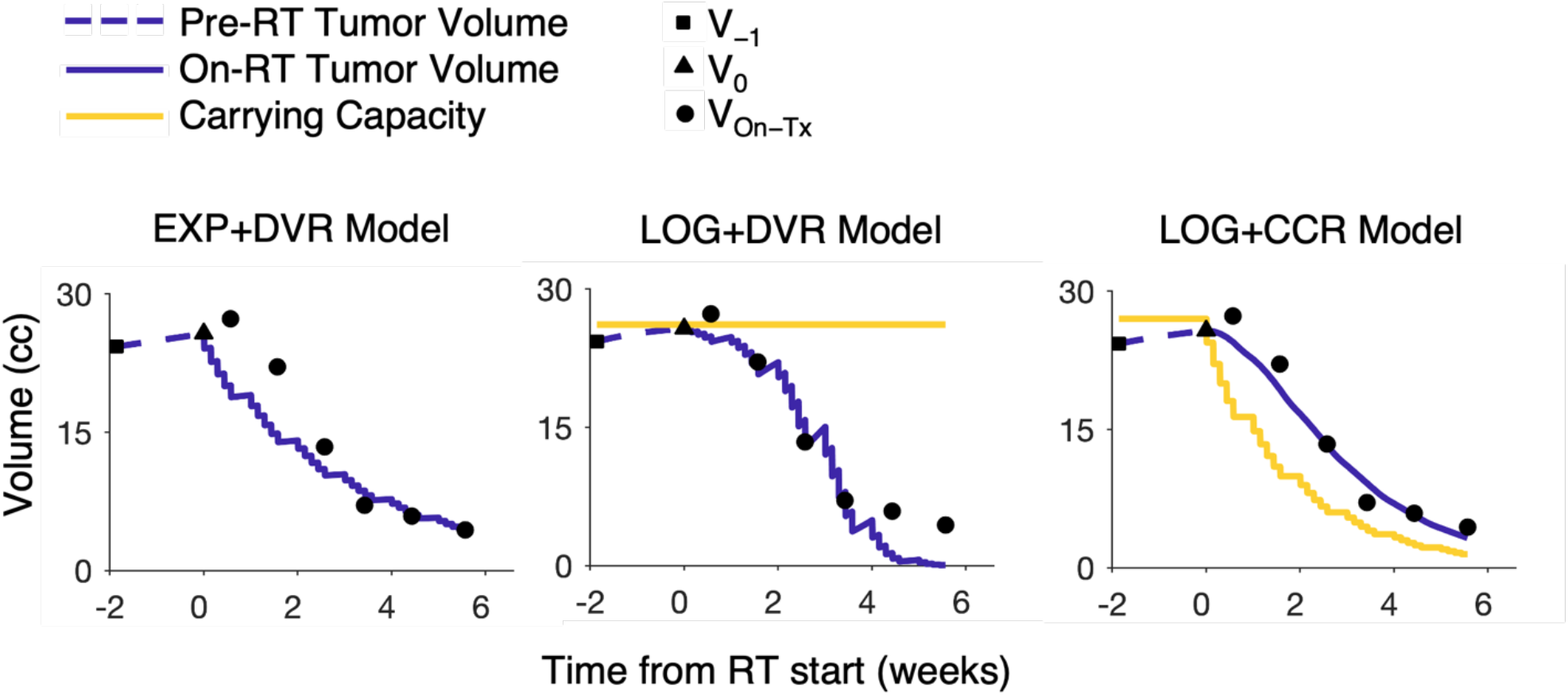
Comparison of example fits of all three models. Model fits to CT-scan derived tumor volume data of one head and neck cancer patient treated at Moffitt Cancer Center. Volume measurements are at time of radiation planning (squares), prior to start of radiotherapy (triangles) and weekly measurements following onset of radiotherapy (circles). The blue curves represent model simulations prior to RT treatment (dashed) and after onset of RT treatment (solid) for each model. In the LOG+DVR and LOG+CCR models, the carrying capacity (yellow curve) bounds the tumor volume growth, which remains constant in the LOG+DVR model and instantaneously drops with each radiation fraction in the LOG+CCR model.

### 2.3 Model Analysis and Comparison

In order to compare the three models, three major types of analysis were performed: (1) parameter sensitivity analysis, (2) parameter identifiability analysis, and (3) analysis of the rates of change in tumor volume.

#### 2.3.1 Parameter Sensitivity Analysis

Sensitivity analysis is used to analyze how the model output changes in response to small perturbations in model parameters. The sensitivity is determined by taking the partial derivative of the model output with respect to the model parameter, given by the sensitivity matrix 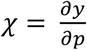 where *p* is the set of model parameters and *y* is the model output. In this case, the model output is tumor volume *V*, with the model parameters for each model listed in Table 1. The sensitivities are then ranked by taking the norm of the sensitivity matrix.(Banks and Tran 2009)

We analyzed the sensitivity of each model parameter along specific ranges. That is, *λ* ∈ (0.055, 0.33) day^-1^, *α* ∈ (0.05, 0.65) Gy^-1^, PSI ∈ (0.1, 0.99), and δ ∈ (0.05,0.75). The lower bound for *λ* was set as one cell division per day (ln 2) multiplied by a 92% cell loss factor with one cell division per day, and the upper bound was set to six times the lower bound to make sure it included previously optimized values for λ(Matsu-ura et al. 2016; Zahid et al. 2021b). The bounds for α were based on a slightly expanded range of α values used to fit response data from a head and neck cancer cohort so as to have a realistic order of magnitude, but still capture more radiosensitive tumors.(Zahid et al. 2021a) We set 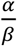 to 10 Gy and did not consider the sensitivity of *β*. The parameter sensitivities were computed and compared for each combination of parameters along the specified ranges.

#### 2.3.2 Parameter Identifiability Analysis

Following the sensitivity analysis, identifiability analysis was used to investigate correlations between parameter pairs. The structural correlation method uses the correlation matrix defined as *C* = *χ*^T^*χ* to compute correlation coefficients *c_ij_* between parameters *i* and *j* (Olufsen and Ottesen 2013). If |*c_ij_*| > *ξ* for *ξ* → 1, then the parameters are correlated and thus, unidentifiable. After removing the least sensitive parameter, this process is repeated iteratively until the parameter set is free of correlations. We used this analysis to determine identifiable parameter spaces for each model along the given ranges.

With the exception of 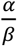 set to 10 Gy and dose set to 2 Gy, a range of values for the remaining parameters were tested in each respective model. The parameter *α* ∈ (0.05, 0.65) Gy^-1^ with intervals of 0.025 Gy^-1^ and the growth rate *λ* ∈ (0.055 to 0.33) day^-1^ with intervals of 0.025 day^-1^. The parameter *δ* ∈ (0.05,0.75) with intervals of 0.025. The upper range of *δ* was set to 0.75 to prevent significantly large changes in the carrying capacity, which would result in unrealistic tumor volume changes. In the EXP+DVR model, the volume at time of radiotherapy planning, *V*_−1_, was set to 60 cc. Since PSI is a parameter rather than K, the equation for K_0_ is rewritten as 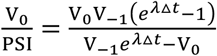. For the LOG+DVR and LOG+CCR models, *V*_0_ was set to 100 cc. For a given 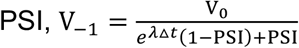. This allows for the tumors to be a similar size at the onset of radiotherapy and differ in volume at the time of radiation planning.

#### 2.3.3 Impact of Parameter Sensitivity on Tumor Volume Trajectories

We first analyzed the impact of the sensitivity of each initial parameter on tumor volume trajectories. Following the identification of identifiable parameters for each model, a MATLAB code constructs a sensitivity matrix where the sensitivity of the model with respect to each parameter is numerically calculated via a forward difference approximation for an input of parameter values. The sensitivity of all parameters over time is then ranked from most sensitive to least sensitive using the two-norm.

For 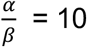 Gy and a dose of 2 Gy, we varied the identifiable parameters for each model. The ranges for parameter values *λ, α*, and *δ* were similar to the identifiable parameters determined previously. PSI ∈ (0.1, 0.99) with intervals of 0.05. Again, *V*_−1_, was set to 60 cc in the EXP+DVR model and *V*_1_ was set to 100 cc in the LOG+DVR and LOG+CCR model. For each model parameter, the parameter sensitivity analysis values were normalized by dividing each parameter’s sensitivity by the largest sensitivity value for the respective parameter. Next, we examined the tumor volume trajectories in the different regions of high, medium, and low sensitivity in the parameter sensitivity of λ. This was done by sampling from circular or spherical regions (in the respective two-dimensional or three-dimensional parameter spaces) of varying sensitivity areas with radii of 0.025 for *λ* in the 3D parameter sensitivity figures.

#### 2.3.4 Maximum reduction of tumor volume (MRV) analysis

The impact of each radiation fraction on tumor volume is mathematically represented by either an instantaneous reduction in tumor volume (EXP+DVR and LOG+DVR model) or an instantaneous reduction in tumor carrying capacity (LOG+CCR model). Since each model has distinct volume dynamics, 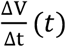 for a given parameter set is unique for each model. To quantify the impact of each radiation fraction on the tumor volume, we introduce the metric of the maximum reduction of tumor volume (MRV), which we define as the largest reduction in tumor volume occurring between any two radiation fractions over the RT treatment schedule.

First, we investigated the impact of the initial tumor state on when MRV occurred, with *λ* = 0.11 day^-1^, *α* = 0.1 Gy^-1^, and δ = 0.2. In order to test the initial tumor state, we varied *V*_−1_ for the EXP+DVR model and PSI for the other 2 models. *V*_−1_ ranged between 5 and 100 with spacing of 5 cc. In the LOG+DVR and LOG+CCR models, *V*_1_ was set to 100 cc, Δ *t* was set to two weeks, and PSI ∈ (0.1, 0.99). For a given PSI, *V*_−1_ was calculated by 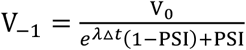. This allowed for a fixed tumor size at the start of RT with different proliferative fractions.

Next, we examined how MRV varies throughout the respective parameter spaces for each model. Here we tested *λ* ∈ (0.055, 0.33) day^-1^, *α* ∈ (0.05, 0.65) Gy^-1^, PSI ∈ (0.1, 0.99), and δ ∈ (0,0.75). For each simulation with its distinct parameter values, we simulated the entire tumor volume trajectory and determined the MRV value and the radiation fraction number at which MRV is achieved.

## 3. Results

### 3.1 Parameter Sensitivity Analysis

In the EXP+DVR model, both *λ* and *α* have similar sensitivity trends (Fig. 2a). in which the sensitivity of both parameters is greatest at low *α* and high *λ* values. For the LOG+DVR model, the sensitivity trends are compared in the 3-dimensional parameter space. There is a distinct region of highest sensitivity for λ at low α and λ values (Fig. 2b). The yellow region demonstrating highest *λ* sensitivity shifts with higher *λ* values and expands to include a larger range of *α* and PSI values. A similar trend is seen for the sensitivity of *α*, however, in this case the region of highest sensitivity does not expand to include all PSI values. The range of values for which PSI is most sensitive is narrower compared to the rest of the parameters – PSI is most sensitive at low *α* values, low PSI values and mid to high *λ* values. For the LOG+CCR model (Fig. 2c), *λ, δ*, and PSI are most sensitive at low δ values and low PSI values. The sensitivities appear less dependent on *λ*. High sensitivity areas for *λ* span across larger PSI values in comparison to *δ* and PSI.

**Figure 2.**
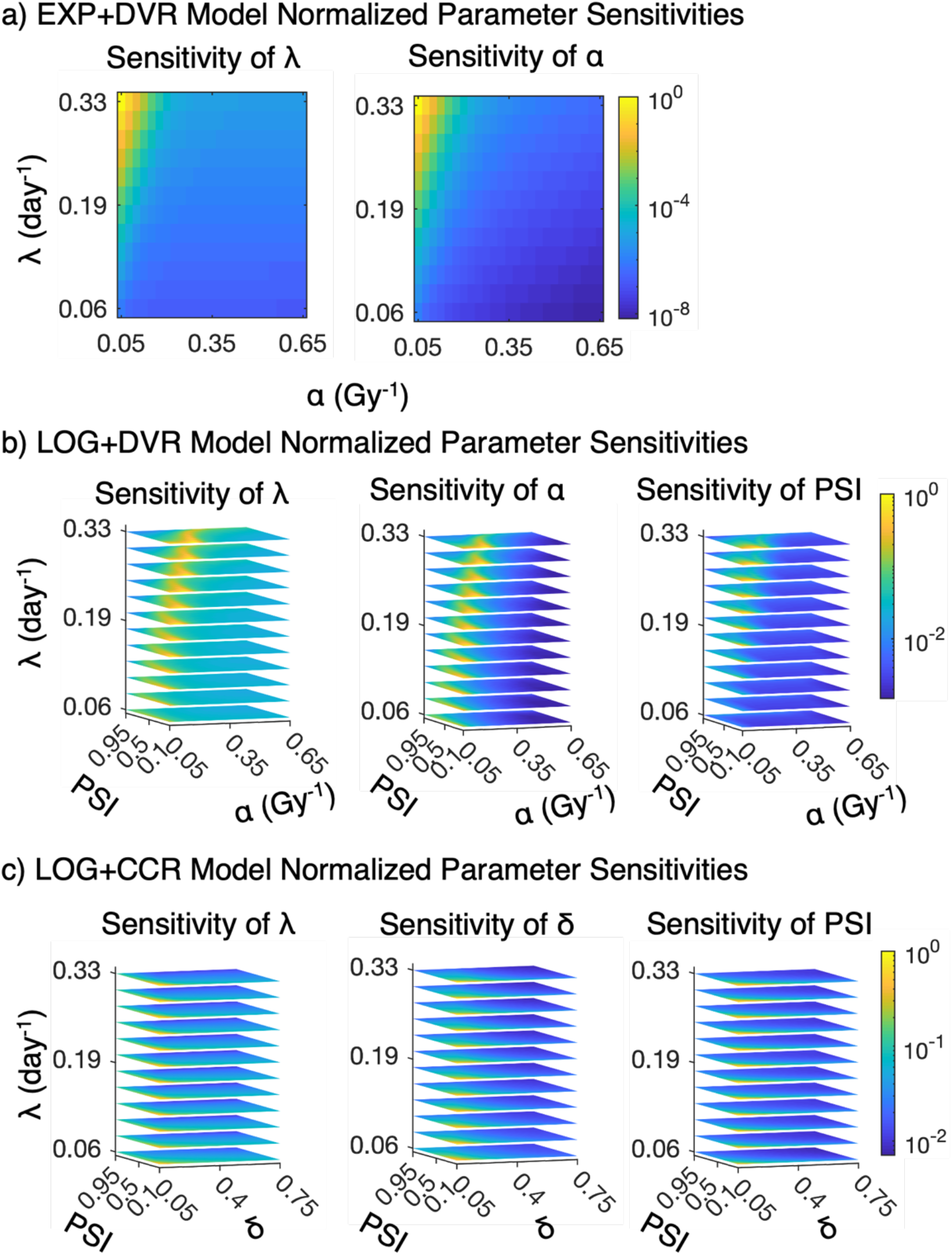
Maps of normalized parameter sensitivity values. 2D and 3D heatmaps show the sensitivity of each parameter for all three models, across their respective parameter spaces. Each parameter sensitivity plot is normalized with its largest parameter value. The color bar quantifies the sensitivity of each parameter for a given input of parameter values.

### 3.2 Parameter Identifiability Analysis

In the EXP+DVR model, λ was always an identifiable parameter, while α is nonidentifiable at low α values. As *λ* increases, α becomes identifiable (Fig. 3a). In the LOG+DVR model, *λ* is always identifiable while α and PSI are identifiable dependent upon the other parameter values (Fig. 3b). The parameter space is divided into 4 types of regions: (1) a region with high α values and low PSI values, where only λ and α identifiable; (2) a region on the other extreme with high PSI values and low α values, where all three parameters are identifiable (except for extremely low λ values, where this region spans all α values); (3) a region where only λ is identifiable, that starts out at low α values when λ is low and drifts to higher α values as λ increases; (4) a few discontinuous regions where only λ and PSI are identifiable: a region at high α and high PSI and a region at low α and low PSI that disappears as λ decreases. In the LOG+CCR model there are three distinct regions: (1) a region with high λ and PSI values where all three parameters are identifiable; (2) a region where only λ and PSI are identifiable, at low λ and high PSI; and (3) the largest region where only λ and δ are identifiable, which primarily has lower PSI and λ values.

**Figure 3.**
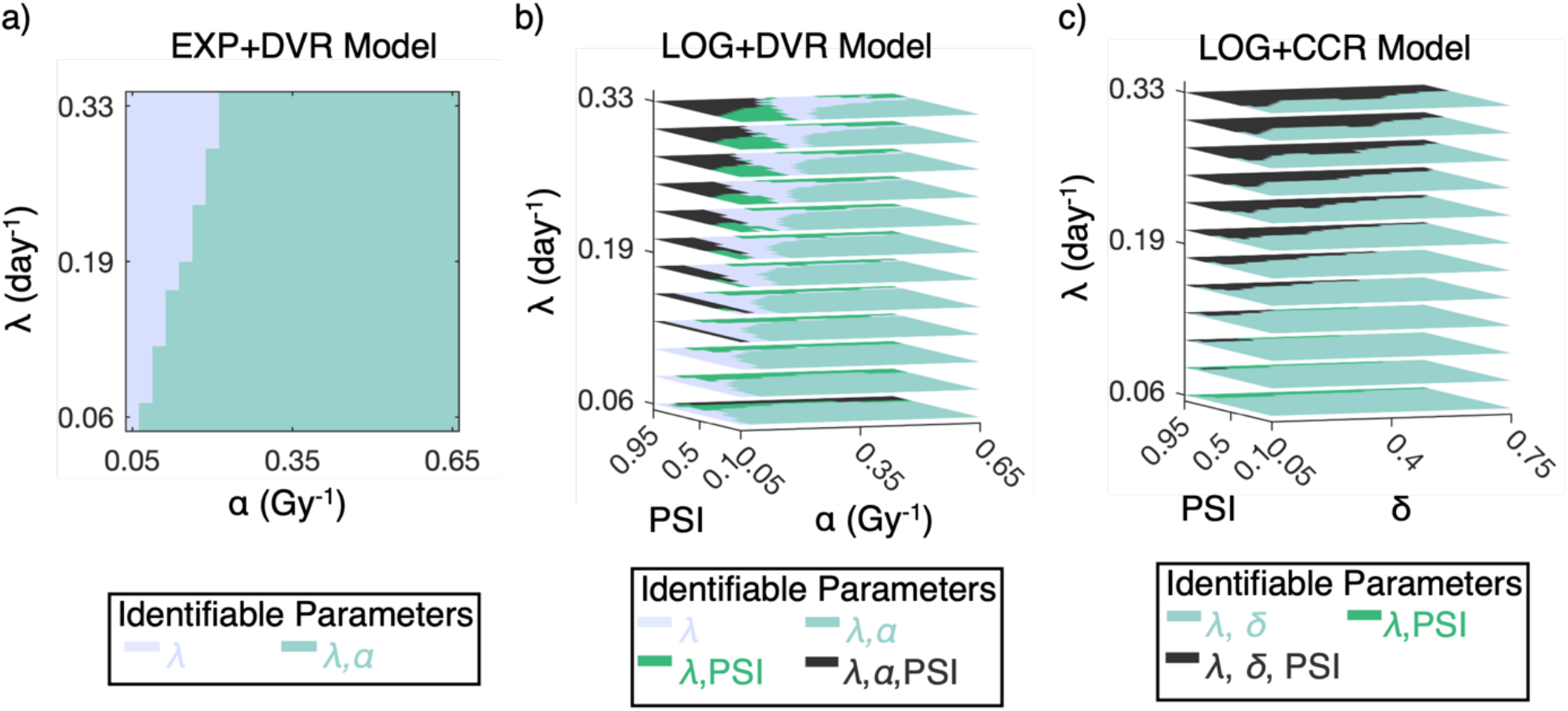
Maps of parameter identifiability regions. Parameter identifiability regions for different parameter values for **a** the EXP+DVR model, **b** the LOG+DVR model, **c** and the LOG+CCR model.

### 3.3 Impact of Parameter Sensitivity on Tumor Volume Trajectories

To further explore the behavior of the models underlying the different degrees of parameter sensitivity, we examined the tumor volume trajectories in the different regions of high, medium, and low sensitivity in the parameter sensitivity of λ, as detailed in the materials and methods (Figure 4A). The tumor volume trajectories for points within each region are depicted for 150 samples per region with a curve to represent the mean and the surrounding shaded area marks volume dynamics one standard deviation from the mean (Fig. 4b). Each point within the regions has a unique value for each identifiable parameter. The ranges for each parameter in each region selected are listed in Supplementary Table 1.

**Figure 4.**
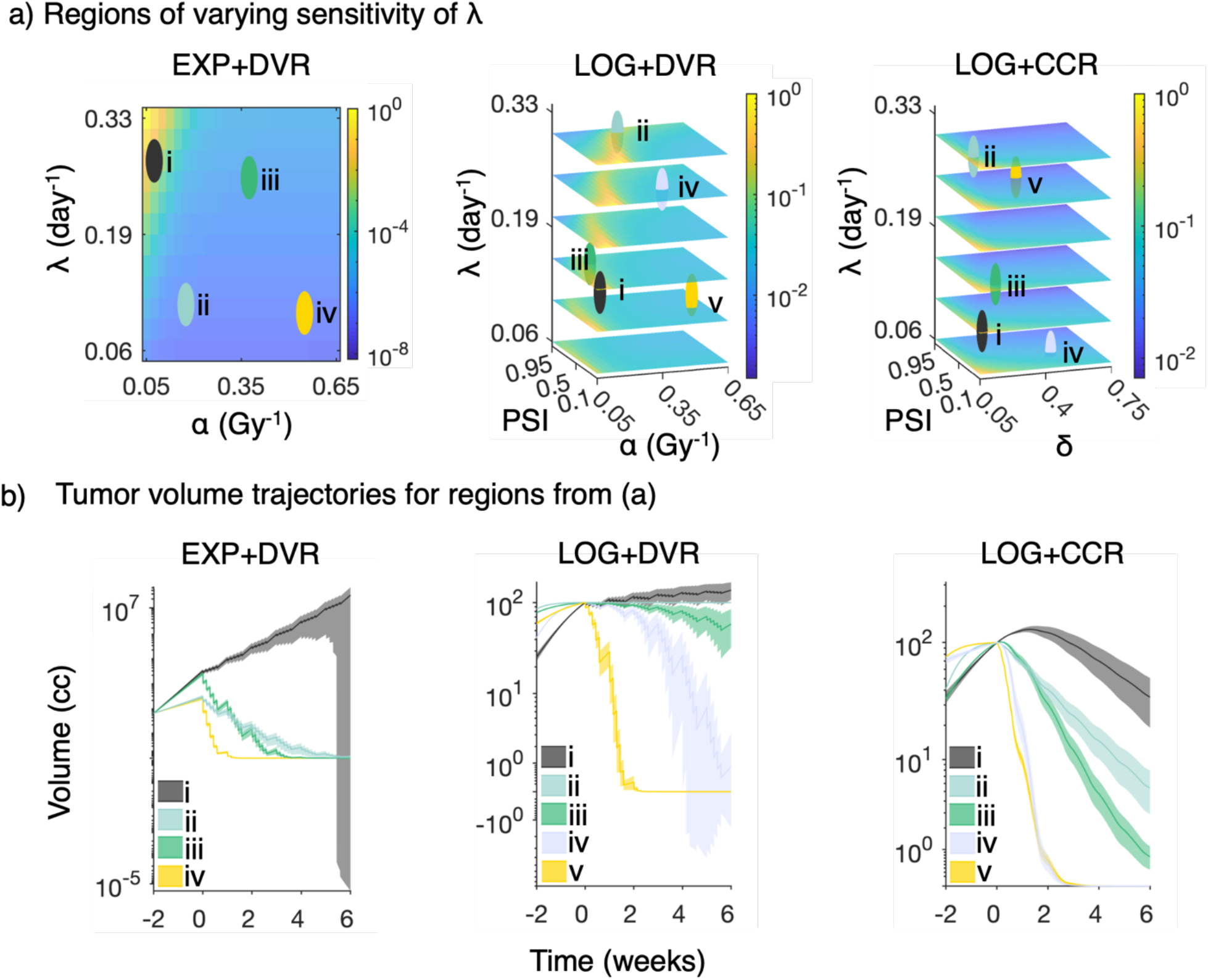
Tumor volume trajectory plots of representative regions of varying sensitivity of λ. **a** 2D and 3D heatmaps quantify the sensitivity of λ defined by the color bar for varying identifiable parameter values. For the EXP+DVR model, each circle encloses a region of *λ* sensitivities defined by a (*α, λ*) pair. For the LOG+DVR and LOG+CCR models, each sphere encloses a region of λ sensitivities defined by a (*λ, α*, PSI) and a (*λ, δ*, PSI) set, respectively. The radii for circles and spheres are 0.025 units. **b** Each curve corresponds to the mean of 150 trajectories of tumor volumes simulated from the parameter values in the corresponding parameter sampling region. Each curve is shaded to visualize trajectories one standard deviation from the mean.

In the EXP+DVR model, in the region where λ is most sensitive (region i), the tumor growth outpaces volume reduction by RT, resulting in net growth. Notably, standard deviation for this region spans a large range of volume dynamics, explaining the high sensitivity of *λ* in this region. The volume versus time data for all points within the region are right skewed after the fourth week of treatment resulting in a standard deviation that is above the mean, which manifests as the dramatic drop in volume for the lower bound of the standard deviation in curve i. For regions ii-iv, volume reduction by RT outpaces the tumor growth rate. Within these regions, as the sensitivity of λ decreases, the tumor volumes generally decrease more rapidly. In the LOG+DVR model, the region with highest sensitivity (region i) again has net tumor growth. The region of medium sensitivity with high PSI values (region ii) has minimal net tumor volume reduction and very little inter-trajectory variability. The volume dynamics for a high PSI tumor will be similar as tumor growth and cell death will occur slowly, if at all, which is why we didn’t test any other regions of high PSI. Regions of low *λ* sensitivity (regions iii-v) include parameter values in which *α* is low and PSI is high and regions where *α* is high and spans all PSI values. As the sensitivity of *λ* decreases across regions, the tumor volume trajectories decrease in size more rapidly. In the LOG+CCR model, a highly sensitive *λ* (region i) occurs when the tumor is highly proliferative (low PSI) and is less radiosensitive (low *δ*). Similar to the other models, as the sensitivity of *λ* decreases in regions ii-v, the rate of tumor volume reduction increases, resulting from increasing *δ* and PSI values. As the sensitivity of *λ* decreases across regions, the tumor volume trajectories vary less from each other.

### 3.4 Maximum reduction of tumor volume (MRV) analysis

For the example simulations fitted to patient data, both the EXP+DVR and LOG+DVR models exhibit tumor growth phases occurring in between radiation fractions and instantaneous tumor cell death at the time of each radiation fraction (Figure 5a). The LOG+CCR model demonstrates more continuous changes in tumor volume as the impact of radiation is not modeled as an instantaneous volume reduction. For each simulation, the MRV magnitude and time to MRV vary. The MRV occurs during the first radiation fraction for the EXP+DVR model, between weeks three and four of RT for the LOG+DVR model, and between weeks one and two of RT for the LOG+CCR model (Figure 5a).

**Figure 5.**
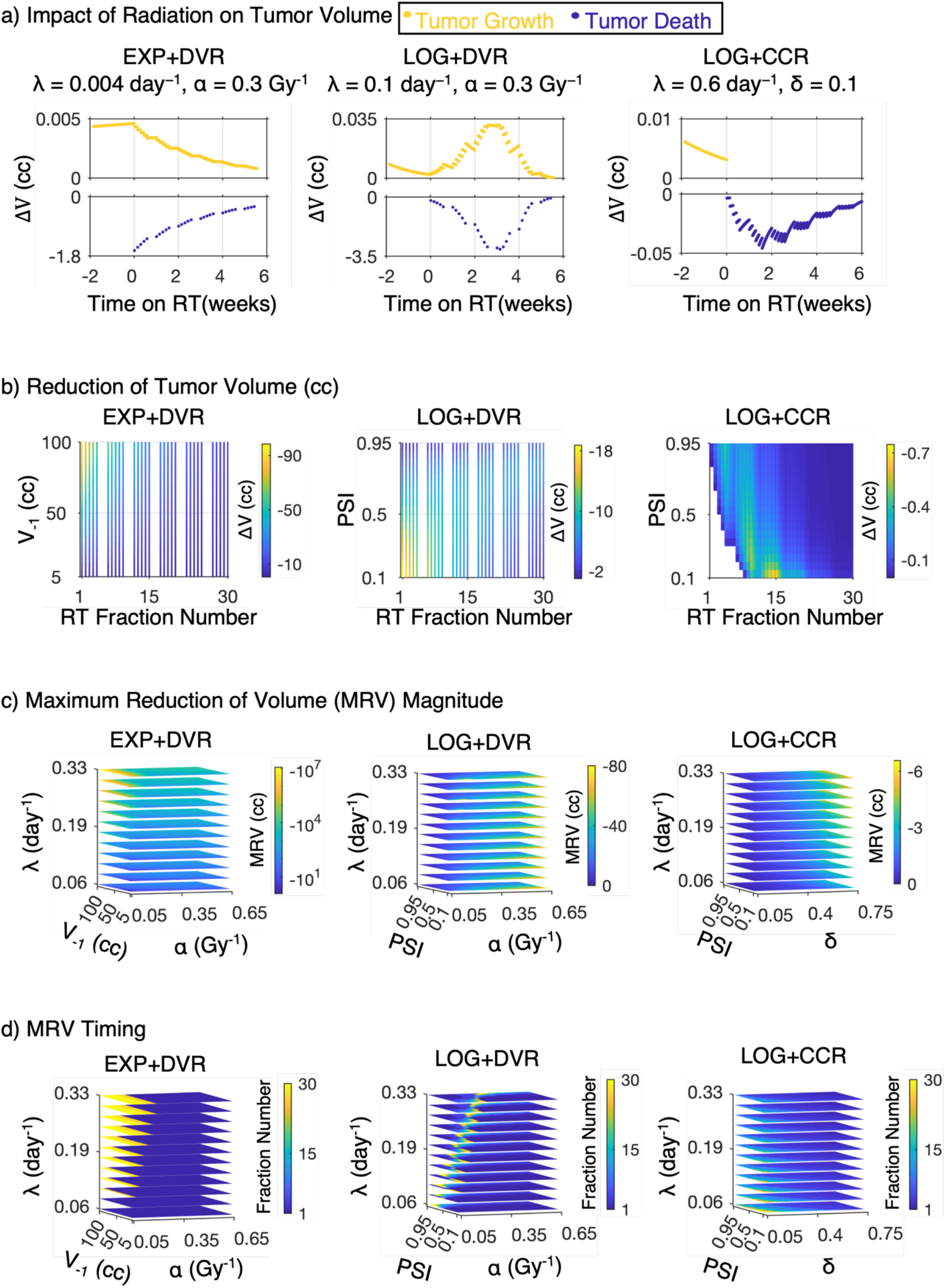
Radiation-induced maximum reduction of tumor volume. **a** The instantaneous change in tumor volumes versus time for the simulated trajectories from Fig. 1b is divided into tumor growth prior to and between radiation fractions (yellow points) and instantaneous tumor volume reduction (blue points) for the EXP+DVR and LOG+DVR model. In the LOG+CCR model, the tumor grows (yellow points) if the tumor is below the carrying capacity and continuously decreases (blue points) in volume, if above the carrying capacity. **b** Heat maps of reduction in tumor volume for the full range of initial conditions: V_−1_ for the EXP+DVR model or PSI for the LOG models over the six-week radiation treatment schedule. Each color quantifies the tumor volume reduction with the associated color bar. In the DVR models, volume reduction only occurs instantaneously at time of radiation fraction. In the LOG+CCR model, tumor volume reduction occurs continuously. **c** Magnitude of maximum tumor volume reduction (MRV) plotted over the respective parameter spaces for all three models as indicated by the color bars. **d** The timing of MRV in radiation fraction number as indicated by the associated color bars. Notably, the maximum number of fractions was 30 fractions, so in these cases the timing of MRV indicates the end of the simulation time.

Changing the initial biological state of the tumor affects the time of MRV differently for the three models (Figure 2b). In the EXP+DVR model, MRV is always at the first radiation fraction with diminishing tumor volume reduction with each subsequent radiation fraction, across all initial tumor volumes. Additionally, tumors with a large *V*_−1_ have a larger magnitude MRV. In the LOG+DVR model, the relative time of MRV is determined by PSI. At low PSI values (PSI < 0.55), tumor volume reduction is greatest with early radiation fractions. Within a given week, tumor volume reductions are greatest on Mondays. Subsequent radiation fractions within the week result in smaller tumor volume reductions. As PSI increases, tumor volume reduction is largest in the middle of radiation treatment. At large PSI values (PSI ≥ 0.85), tumor volume reduction is largest at the last radiation fraction. In this case, tumor volume reductions are greatest on Fridays in each week of RT. In the LOG+CCR model, MRV always occurs before the fifteenth radiation fraction (end of week 3 of RT). MRV occurs earlier with a high PSI (PSI > 0.5). For low PSI values (PSI ≤ 0.5), the MRV occurred at or prior to the ninth RT fraction (middle of week 2 of RT). After MRV is reached, the reduction of tumor volume decreases in magnitude with each subsequent RT treatment week.

The timing and magnitude of MRV also varies across the respective parameter spaces for each model (Figure 5c-d). In the EXP+DVR model, the values of MRV range from −1.20 to −1.16 × 10^7^ cc. MRV has the greatest magnitude at high *λ* and low *α* values, and in these cases, the time to MRV occurs at the end of the treatment course at radiation fraction 30 (Figure 5d). These are cases in which the tumor growth outpaced volume reduction from radiation and continues to grow. For most input values in which radiation induces volume reduction, MRV occurs with the first radiation fraction in the radiation treatment course.

In the LOG+DVR model, the MRV value ranges from −0.49 to -78.91 cc (Figure 5C). MRV values are greatest in magnitude when PSI is low and *α* is high. This occurs when the tumor’s proliferative capacity is high and the cell death parameter is large, resulting in sharp volume reductions with each radiation fraction. As PSI increases, the MRV decreases – this is most apparent at high α values. Although regions of low PSI and large *α* values exhibit large absolute MRV values (the largest MRV of -78.91 cc occurs at a *λ* = 0.33 day^-1^, *α* = 0.18 Gy^-1^, and PSI = 0.01), the tumor volume continues to grow more rapidly than the radiation induced volume reductions. The number fractions to MRV is highly parameter-dependent with the region of longest time to MRV drifting expanding from a small subregions exclusively at higher PSI values to two subregions at both high and low PSI values, as λ increases, even though the vast majority of parameter combinations have MRV early on the treatment course (Figure 5d).

In the LOG+CCR model, MRV values range from -0.0970 to -6.5673 cc (Figure 5C). MRV values are largest at high *δ* values and high *λ* values. A trend across varying PSI values is not evident. At high *λ* values, high *δ* values, and high PSI values a small number of fractions (<5) are required to reach MRV, while when *λ, δ*, and PSI are all low the number of fractions required to reach MRV increases.

## 4. Discussion

We conducted an extensive comparison of three different models of tumor volume response to RT. The parameter identifiability and sensitivity analyses revealed the interdependence of the different model parameters and may inform parameter calibration in any further usage of these models. We further examined the impacts of the parameter sensitivity on the tumor volume trajectories to obtain further understanding of model behavior. Finally, we examined the changes in *ΔV*/*Δt* (*t*) to elucidate the implications of model selection.

The sensitivity analyses demonstrated that all parameter sensitivities are highly dependent on the nominal values of the other parameters. Additionally, the regions of highest parameter sensitivity are where tumor growth outpaces tumor volume reduction by RT, and as RT efficacy increases parameter sensitivity values decrease. This decreasing sensitivity of *λ* with high PSI values may explain why setting *λ* uniform across a patient cohort did not affect goodness of fit significantly (Zahid et al. 2021b). It is also important to note we selected a wide range of parameter values to assure that any results at extreme values would also be observed, although many of these values may not be physiologically observed. Therefore, parameter ranges in this analysis are selected for the sake of thorough model analysis and should not be considered as calibrated ranges.

For all three models, there are large regions of the parameter spaces where λ and the respective radiosensitivity parameter (α for DVR models and δ for the CCR model) are identifiable, which is important for being able to separately parameterize tumor growth and RT response. Notably, these regions of the parameter space require sufficiently large tumor growth rates. For both models with logistic growth, there are sizeable parameter regions where PSI is identifiable but the radiosensitivity parameter is not. However, there are additional regions where PSI is identifiable, while the radiosensitivity parameter is not. This may indicate that with more complex growth models that attempt to encode more biological detail, it may be difficult to distinguish between the effects of RT and changes in tumor growth properties (e.g. proliferative fractions), in intermediate parameter regions.

Our newly presented MRV analysis, allows for functional analysis of the tumor volume dynamics during the course of treatment across all three models. For the EXP + DVR model, fast growing tumors can outgrow tumor volume reduction due to radiation in this model resulting in largest instantaneous MRV occurring at the end of the treatment course. On the other hand, smaller, slower growing, more highly radiosensitive tumors have a smaller MRV values, which occur with the first radiation fraction. For the LOG+DVR model, in the high *λ* sensitivity region, the time to MRV occurs at later radiation fractions. This is in contrast to regions where the parameters are less sensitive and *α* is identifiable, where MRV occurs in the early radiation fractions. Finally, in the LOG +CCR model, MRV values are relatively small and generally occur towards the beginning of the treatment course. MRV values are largest in the EXP+DVR model and orders of magnitude smaller in the LOG+CCR model. The range for MRV values differs drastically across each model. The maximum MRV (most negative) value for the EXP+DVR model is more than six magnitudes larger than the maximum MRV for the LOG+DVR model. On the other hand, the maximum MRV for the LOG+CCR model is ten times smaller than the LOG+DVR model. This is due to the fact that after the carrying capacity drops below the tumor volume, the tumor continuously decreases, allowing for the smaller instantaneous volume reductions that add up over tie to achieve the same net volume reduction as seen in the DVR models, despite the significantly larger instantaneous volume reductions.

The unique MRV analysis and subsequent results bring up some interesting clinical questions, such as: Is there a point of diminishing returns for RT? Is there truly regrowth of the tumor in between radiation fractions? The results of this analysis generate hypotheses that can be validated by comparison of tumor volume measurements over time. These and other questions can be tested with appropriate pre-clinical or clinical data, and the results of these tests may inform us which aspects of the different models are most representative of the true underlying biology, which may inform further model refinement. Some of the results may require more granular measurements than are typically collected in the clinical setting, but hopefully such data collection would be motivated by the clinical utility of the models. Ultimately, more detailed study of these models may enable more appropriate model selection on a disease-site or patient-by-patient basis.

## Supplementary Information

**Supplementary Table 1.** Ranges of parameters sampled for the EXP+DVR model for each region on the parameter sensitivity figure for *λ* (Figure 4).

| EXP+DVR Model |  |  |
| --- | --- | --- |
| Region | Range of $\alpha$ (Gy<sup>-1</sup>) | Range of $\lambda$ (day<sup>-1</sup>) |
| i | 0.52–0.58 | 0.08-0.13 |
| ii | 0.05-0.10 | 0.23-0.31 |
| iii | 0.35-0.40 | 0.24-0.29 |
| iv | 0.15-0.20 | 0.09-0.14 |

**Supplementary Table 2.** Ranges of parameters sampled for the LOG+DVR model for each region on the parameter sensitivity figure for *λ* (Figure 4).

| LOG+DVR Model |  |  |  |
| --- | --- | --- | --- |
| Region | Range of $\alpha$ (Gy<sup>-1</sup>) | Range of PSI | Range of $\lambda$ (day<sup>-1</sup>) |
| i | 0.13-0.18 | 0.23-0.28 | 0.23-0.28 |
| ii | 0.05-0.10 | 0.13-0.18 | 0.13-0.18 |
| iii | 0.30-0.35 | 0.83-0.88 | 0.28-0.33 |
| iv | 0.15-0.20 | 0.73-0.78 | 0.13-0.18 |
| v | 0.08-0.13 | 0.88-0.93 | 0.18-0.23 |
| vi | 0.56-0.61 | 0.48-0.53 | 0.10-0.150 |

**Supplementary Table 3.**
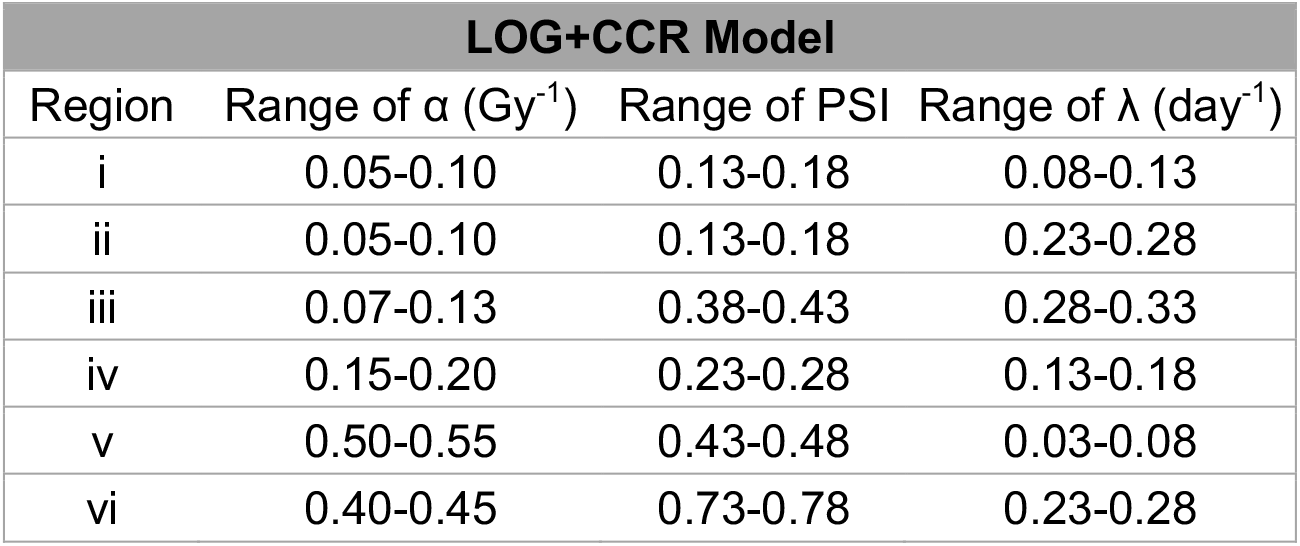
Ranges of parameters sampled for the LOG+CCR model for each region on the parameter sensitivity figure for *λ* (Figure 4).

| LOG+CCR Model |  |  |  |
| --- | --- | --- | --- |
| Region | Range of $\alpha$ (Gy<sup>-1</sup>) | Range of PSI | Range of $\lambda$ (day<sup>-1</sup>) |
| i | 0.05-0.10 | 0.13-0.18 | 0.08-0.13 |
| ii | 0.05-0.10 | 0.13-0.18 | 0.23-0.28 |
| iii | 0.07-0.13 | 0.38-0.43 | 0.28-0.33 |
| iv | 0.15-0.20 | 0.23-0.28 | 0.13-0.18 |
| v | 0.50-0.55 | 0.43-0.48 | 0.03-0.08 |
| vi | 0.40-0.45 | 0.73-0.78 | 0.23-0.28 |

